# Tech Note – Adaptive Sampling: targeted Oxford Nanopore long-read sequencing

**DOI:** 10.1101/2025.08.31.672778

**Authors:** Nick Crang, Orlando Contreras-López, Remi-André Olsen, Fahri Hasby

## Abstract

Oxford Nanopore Technology (ONT) has developed a long-read sequencing method known as “adaptive” or “Read Until” sequencing. Unlike Illumina sequencing, this ONT read data is available in real-time during the run, rather than only being available when the sequencing run is complete and it focuses on a specified region of interest (ROI) within the sequence data. We investigated how this technique compared to standard sequencing under a variety of conditions and found it consistently gave a multifold increase in coverage of the ROI compared to standard sequencing, though a strong positional effect was observed for ROI efficacy.

Adaptive sampling can be used to selectively enrich or deplete sequences from a library in real-time, enhancing coverage of regions of interest without requiring additional library pre-processing

## Introduction

Oxford Nanopore Technology (ONT) has developed a long-read sequencing method known as “adaptive” or “Read Until” sequencing. During typical ONT sequencing, DNA strands are read via changes in electrical potential as the molecules transit through pores in a specially prepared membrane. Unlike Illumina sequencing, this ONT read data is available in real-time during the run, rather than only being available when the sequencing run is complete. In adaptive sampling (AS), the sequence data is both generated and compared in real-time to that of a selected region of interest (ROI) within the genome. If the sequence of a DNA strand does not match the ROI, the voltage is reversed and the strand is ejected from the pore, leaving the pore available to read another strand. The user provides the FASTA sequence of the reference genome along with the coordinates of the ROI as a BED file. Multiple non-overlapping different ROIs can be sampled simultaneously. This technology allows for sample enrichment without the intensive and time-consuming library preparations required by, for example, Cas9 targeted enrichment for Nanopore sequencing. AS increases the proportion of low abundance reads in metagenomic samples (Martin et al. 2022) and has the potential for characterising regions of high variance (Weilguny et al. 2023). Note that although it is possible to use AS to deplete sequences as well as enrich them, this feature was not tested in this study.

This project consisted of five distinct branches of inquiry

1. Does AS give greater overall coverage of ROI’s than standard sequencing?
2. Does the length and/or number of the ROI’s impact the enrichment from AS?
3. Does AS reduce the effective lifespan of the flow cells?
4. Is it possible to read into gaps between ROIs, allowing unknown regions to be read into from known ones?
5. Is the enrichment more effective if the ROI is used as the reference genome?

To answer these questions two separate experiments were performed using two separate flow cells (see Table 1 for a summary of run conditions for both experiments). In one experiment we performed a run using a single flow cell, where half the channels were reserved for standard sequencing and half for adaptive sampling. This was termed the “reserved channel” experiment. In the “reserved channel” experiment we aimed to determine both if adaptive sampling reduced the lifespan of the flow cell and provide a direct comparison of whether adaptive sampling would increase the overall coverage of an ROI relative to standard sequencing. We have observed that both read number and quality varies greatly between flow cells and therefore concluded that the only way to effectively compare AS with standard sequencing was by partitioning a flow cell. Doing so allows us to run AS on half the channels in a flow cell, and standard sequencing in the remainder. This approach removed potential confounding variables from loading, the library itself and the flow cell, when compared to running on two separate flow cells.

**Table 1.** Summary of run conditions and their purpose.

| **Experiment 1 – Reserved channel experiment** |  |  |  |  |
| --- | --- | --- | --- | --- |
| **Run name** | **ROI % genome** | **No. of ROIs** | **Run time / hours** | **Purpose** |
| Channels 1-1499 | 8 | 1 | 72 | AS using a single ROI performed on half the channels in the flow cell |
| Channels 1500-3000 | 0 | 0 | 72 | Standard sequencing performed on half the channels in the flow cell |
| **Experiment 2 – Multiple run experiment** |  |  |  |  |
| P29602_101 | 0 | 0 | 0.5 | Positive control using standard sequencing to confirm flow cell functionality at the start of the experiment |
| P29602_01pct | 0.1 | 1 | 1 | AS using a single ROI to test if length of the ROI effects enrichment |
| P29602_025pct | 0.25 | 1 | 1 |  |
| P29602_05pct | 0.5 | 1 | 1 |  |
| P29602_1pct | 1 | 1 | 1 |  |
| P29602_4pct | 4 | 1 | 1 |  |
| P29602_8pct | 8 | 1 | 1 |  |
| P29602_ROI_as_ref | 1 | 1 | 2 | AS without a BED file, instead using an ROI in place of the reference genome to test an alternative mapping strategy |
| P29602_NoAS | 0 | 0 | 0.5 | Positive control using standard sequencing to confirm flow cell functionality at the halfway point of the experiment |
| P29602_01pct_10ROIs | 1 | 10 | 1 | AS using 10 ROIs, enrichment compared to P29601_1pct |
| P29602_01pct_80ROIs | 8 | 80 | 1 | AS using 80 ROIs, enrichment compared to P29602_8pct |
| P29602_day3_gapfill | 0.5 | 1 | 19 | AS using a reference file containing 500 Kbp of NNN simulating a gap in the genome |

The other experiment used a single flow cell loaded with a single library and was termed the “multiple run” experiment. The sequencing run was stopped and restarted to be able to adjust the sequencing settings. The first run in this experiment did not perform AS and was a positive control to confirm the flow cell functionality (this test was performed at the halfway point of the experiment as well). Subsequently, several AS runs were set up varying the number and the lengths of the ROIs selected to answer questions 1-4.

## Results

AS enrichment was clearly observed in the “reserved channel” experiment shown in Fig. 1. Here 1500 channels were selected to be used for AS and 1500 channels did not perform AS. The single ROI covering 8% of the genome (also used in the “multiple run” experiment) was selected for this test. The channels performing AS yielded 16.96x coverage for the ROI compared with 4.67x with standard sequencing for the same region, an improvement of 12.29x. This is slightly lower than half the maximum 40x coverage reported from ONT ([Community Nanoporetech info sheet](https://community.nanoporetech.com/info_sheets/adaptive-sampling/v/ads_s1016_v1_revj_12nov2020/considerations-for-experimental-design)) but this is to be expected since the flow cell was split between AS and standard sequencing, so only half the total sequencing capacity was used for each. Here 3.63 fold enrichment was seen rather than the 5-10 fold claimed by ONT, though this also may be as a result of the decreased total sequencing compared to a non-split flow cell experiment.

**Fig 1.**
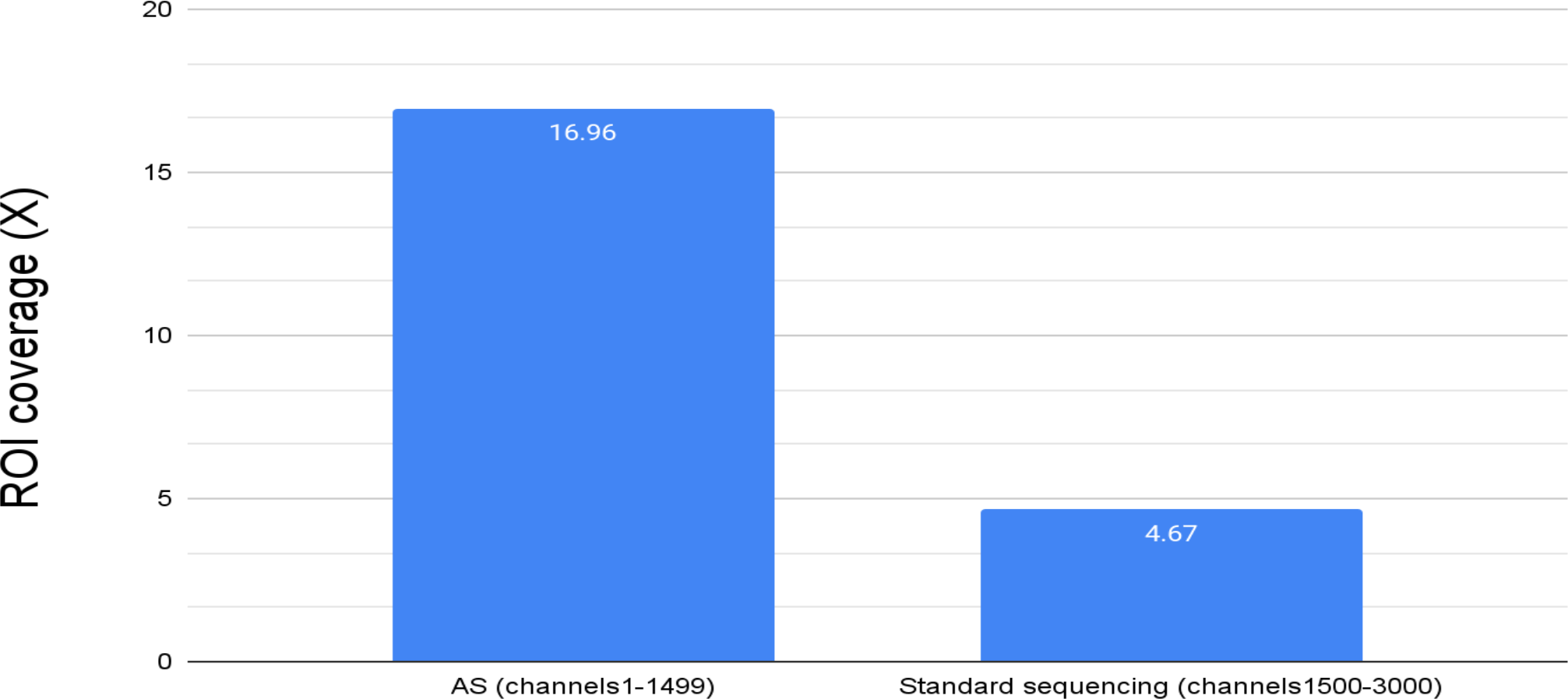
Adaptive sampling increases the coverage of ROIs vs standard sequencing. A flow cell run was split with 1500 channels reserved for adaptive sampling and 1500 performing standard sequencing. The total fold coverage of an ROI comprising 8% of the total genome is displayed here. The total coverage was 12.29 X greater when adaptive sampling was used. AS = adaptive sampling.

While the coverage increased with AS, both the N50 and read length distribution of reads that passed filter was comparable to that seen from standard sequencing (see Appendix 1). It should be noted that there are a far greater number of reads <400bp that do not pass filtering during AS but that is to be expected as any reads rejected for not matching the ROI will be in this size range.

In the “multiple run” experiment we performed multiple short sequencing runs on the same library using a range of single ROIs covering different proportions of the genome. As well as these single ROIs, we also performed runs that covered the same proportion of the genome (1 and 8%) but split over 10 or 80 different ROIs scattered throughout the genome rather than as a single ROI in chromosome 1 or 2. In addition we used single ROIs to test an alternative mapping strategy as well as the ability to sequence into unknown sections of the genome. We observed, somewhat surprisingly, that the size of the ROI did not influence the observed enrichment within the size range tested (0.1-8% of the total genome, Fig 2.). This may seem counterintuitive at first, however, although more sequences are required for 1x coverage of a longer ROI sequence than a shorter one, the sequences mapping to a shorter ROI are rarer. As only sequences that map to the ROI will be sequenced fully during adaptive sampling, when a shorter ROI is used a greater proportion of sequences are discarded before being fully sequenced than when a longer ROI is used.

**Fig 2.**
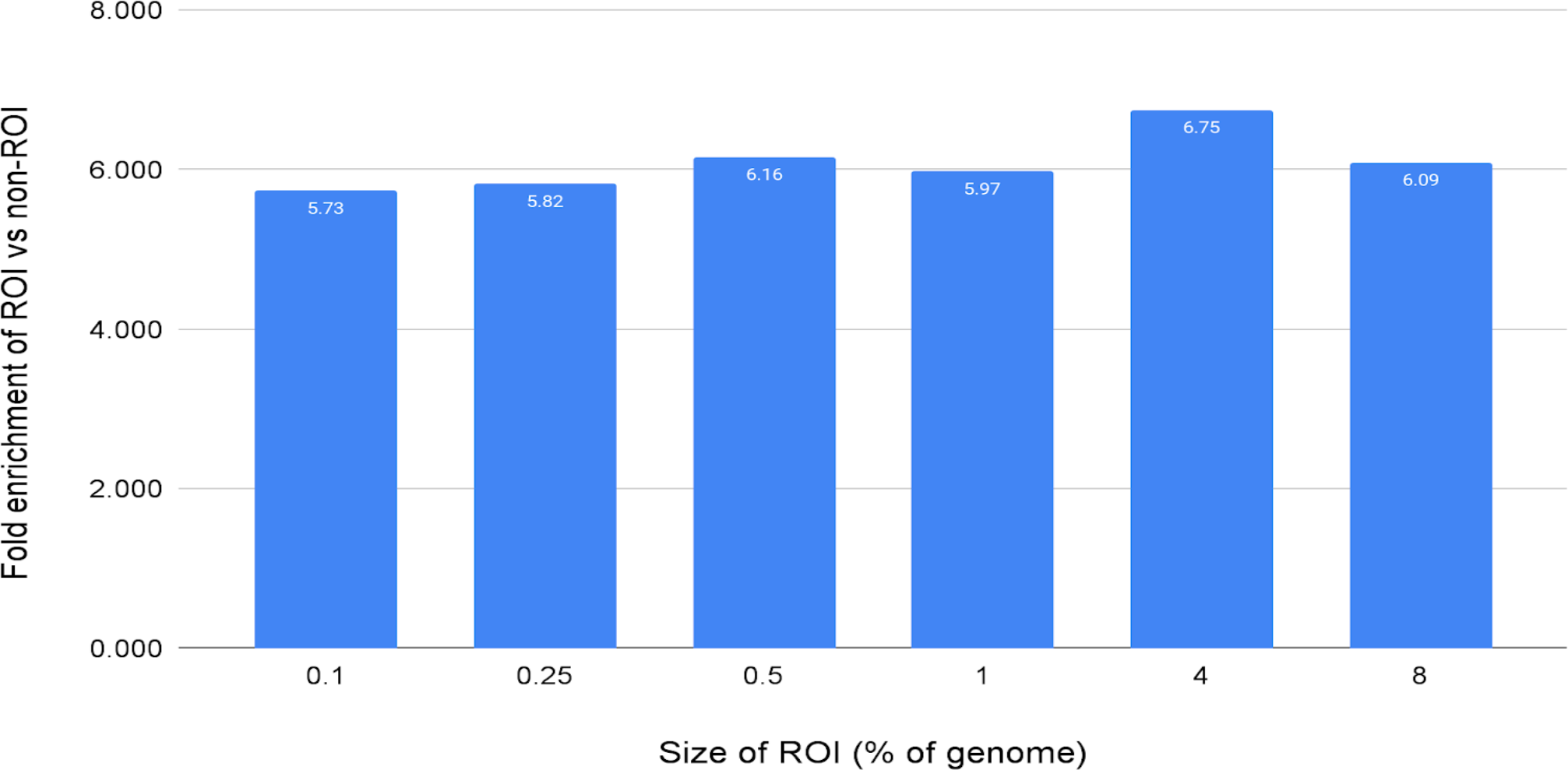
Length of ROI does not affect the relative enrichment of coverage. A single library was sequenced using adaptive sampling using different BED files with varying ROI lengths. The ROIs ranged from 0.1-8% of the total genome in size and each was sequenced for 1 hour. Total coverage varied from 5.7-6.7x and the number of reads from 0.3-0.57 million reads.

The total coverages seen in this experiment were much lower than that observed during the reserved channel flow cell test (5.7-6.7x vs 16.9x). This was expected, since during this experiment the sample library was run multiple times with different BED files for only a single hour each. This gave far fewer total reads per ROI (approx. 0.5 million each) than the split flow cell experiment (5 million reads). It is worth noting that the alternative mapping strategy which used the ROI comprising 1% of the total human genome as the reference genome for mapping gave only 4.2x enrichment compared to the 5.97x enrichment achieved via standard AS mapping of the ROI to the whole genome.

Similarly to the length, we observed that the number of individual sequences making up an ROI did not affect the enrichment obtained (Fig 3.). A single ROI covering 8% of the total genome was compared to 80 ROIs covering 0.1% each, resulting in the same total length of ROI. These resulted in 6.1 and 6.2x enrichment respectively, though there was localised variation seen, with the coverage of some ROIs dropping to 0. This is highlighted in Fig 4.

**Fig 3.**
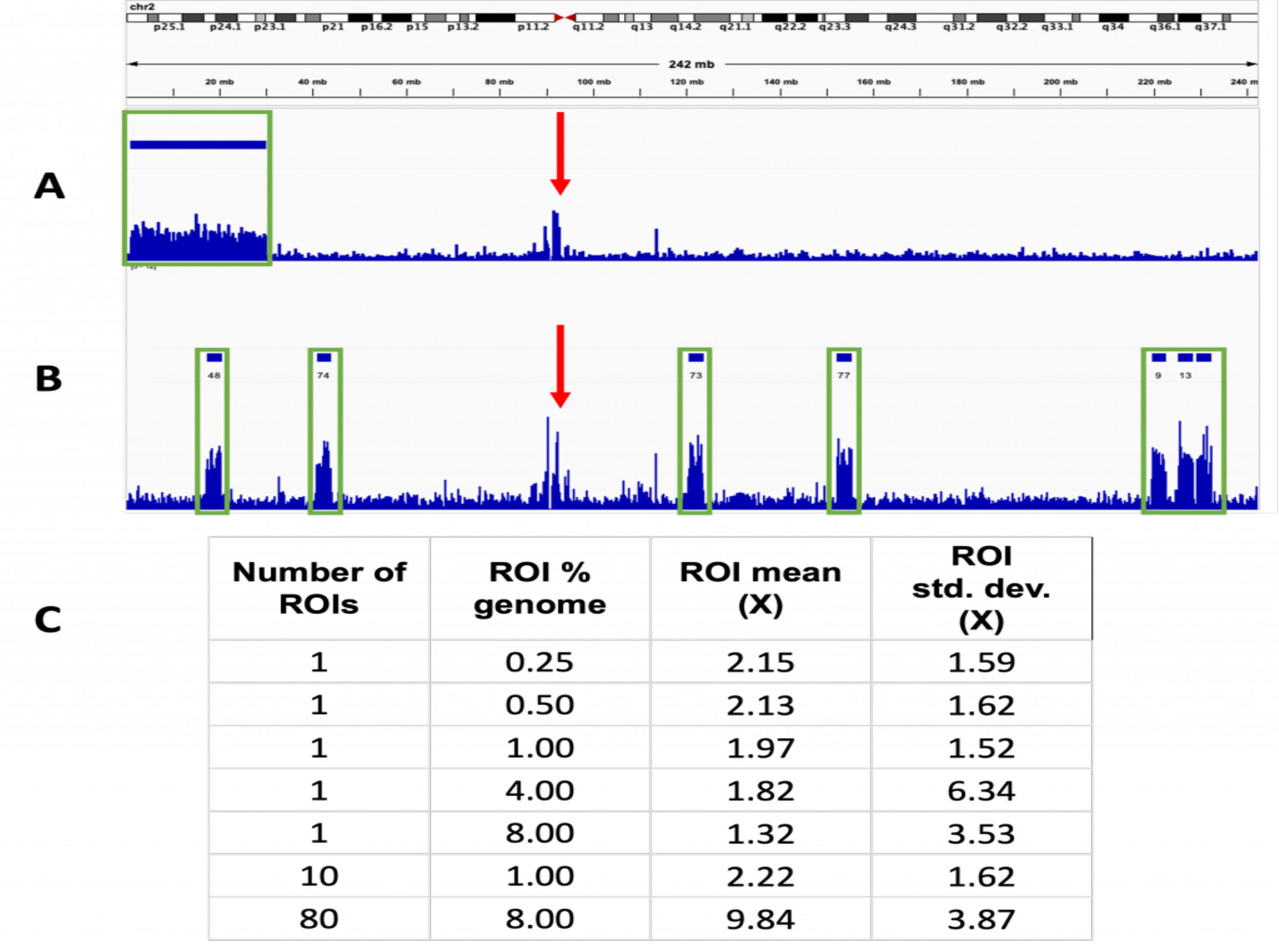
The number of ROIs does not affect the enrichment. Track view taken from chromosome 2 for two separate runs using different numbers of ROIs. Four separate tracks are shown, the top two (A) for a sequencing run using a single ROI and the bottom two (B) for a run using multiple ROIs. The top track indicated the position of the ROI which is displayed as solid bars above the tracks. The track below shows the sequence coverage for the corresponding run with the height of the lines in the lower track indicating the degree of coverage at that point. In both cases, increased coverage was also seen outside of the ROIs at the centromeres. The two red triangles indicate the centromere of the chromosome. Green rectangles indicate positions of enrichment at ROIs and red arrows highlight sequence enrichment at the centromeres. **A** – Used a single ROI spanning chromosomes 1 and 2, measuring 8% of the total genome. Non-specific enrichment also seen around the centromere **B** – Used 80 ROIs each measuring 0.1% of the total genome. **C** – Table summarising the size, number and coverage observed during the sequencing runs with different ROIs.

**Fig 4.**
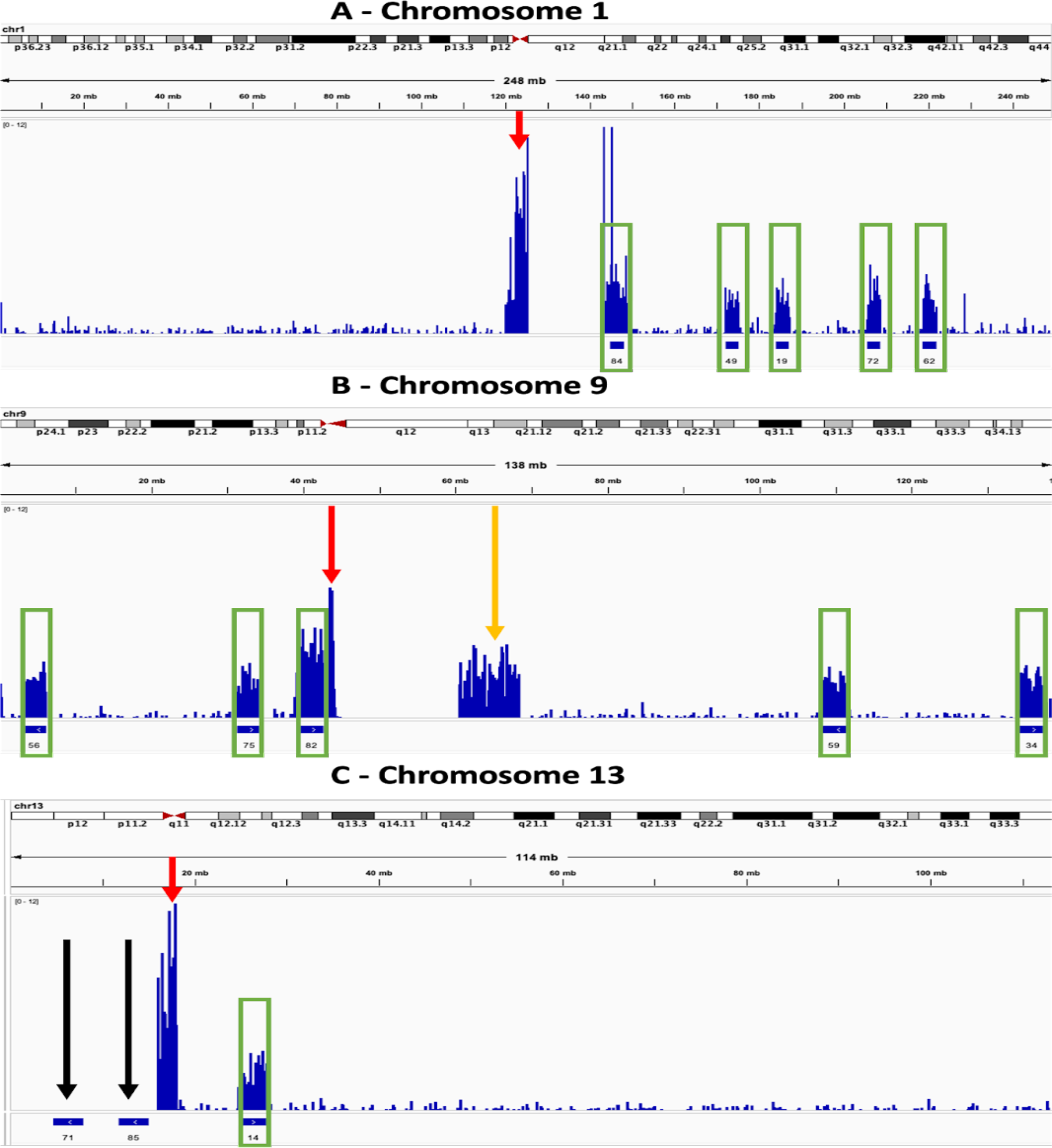
Enrichment varies between ROIs depending on location within the genome. Track views taken from several chromosomes. The upper track for each displays the sequencing coverage, which is indicated by the height of the bar. The lower track indicates the position of ROIs, which are displayed as numbered solid blue bars. Green rectangles further highlight the positions of enrichment at ROIs and red arrows highlight sequence enrichment at the centromeres. **A** – Enrichment is seen at centromeres in addition to the ROIs. **B** – Enrichment is sometimes seen in regions outside of the ROIs and centromeres. Orange arrow highlights enrichment outside the ROIs. **C** – Sometimes enrichment does not occur at specific ROIs. Black arrows indicate the ROIs which were not enriched

In Fig. 4A-C the tendency of reads to be mapped to the repetitive sequences found in the centromeres of the chromosomes is clearly displayed, with the total amount of reads being assigned to them often being considerably higher than that seen from the ROIs. The need to consider the architecture of the genome when designing ROIs is highlighted in Fig. 4B, where high coverage is seen in q13-21 of chromosome 9, a region well outside any of the ROIs. This can be explained by the fact that q13-21 shares considerable homology with p12 (Starke et al. 2002), so reads from the ROI can be mapped to both positions. The failure of reads to map at all to certain ROIs is shown in Fig. 4C, where two of the three ROIs in chromosome 13 had no sequences mapped to them at all, despite average levels of enrichment seen for the third ROI. The reason for this is unclear as there is nothing annotated from p11.2-p13, the region spanned by the two ROIs, however it also had no coverage from standard sequencing. This suggests that the region is generally intractable to sequencing and/or mapping rather than this being a problem specific for AS itself. Similar such regions should be avoided in future when designing ROIs.

We tested the ability of AS to read into unknown sequences. An ROI with a 500 kbp “unknown region” where the sequence was replaced with Ns was constructed in hg38 chr1 and used for AS. We see (Fig. 5) that this does occur during AS with reads being anchored by the flanking regions continuing sequencing and providing some coverage into the unknown region. The efficacy of this is limited by the relatively short read lengths and coverage drops dramatically compared to the surrounding region. In this instance we used an assembly gap significantly larger than the N50 read size (11.78 Kb estimated by MinKNOW). The nature of ONT sequencing lends some intriguing possibilities to this application, as hypervariable regions could be accurately read and assembled in a way not possible with short read shotgun sequencing.

**Fig 5.**
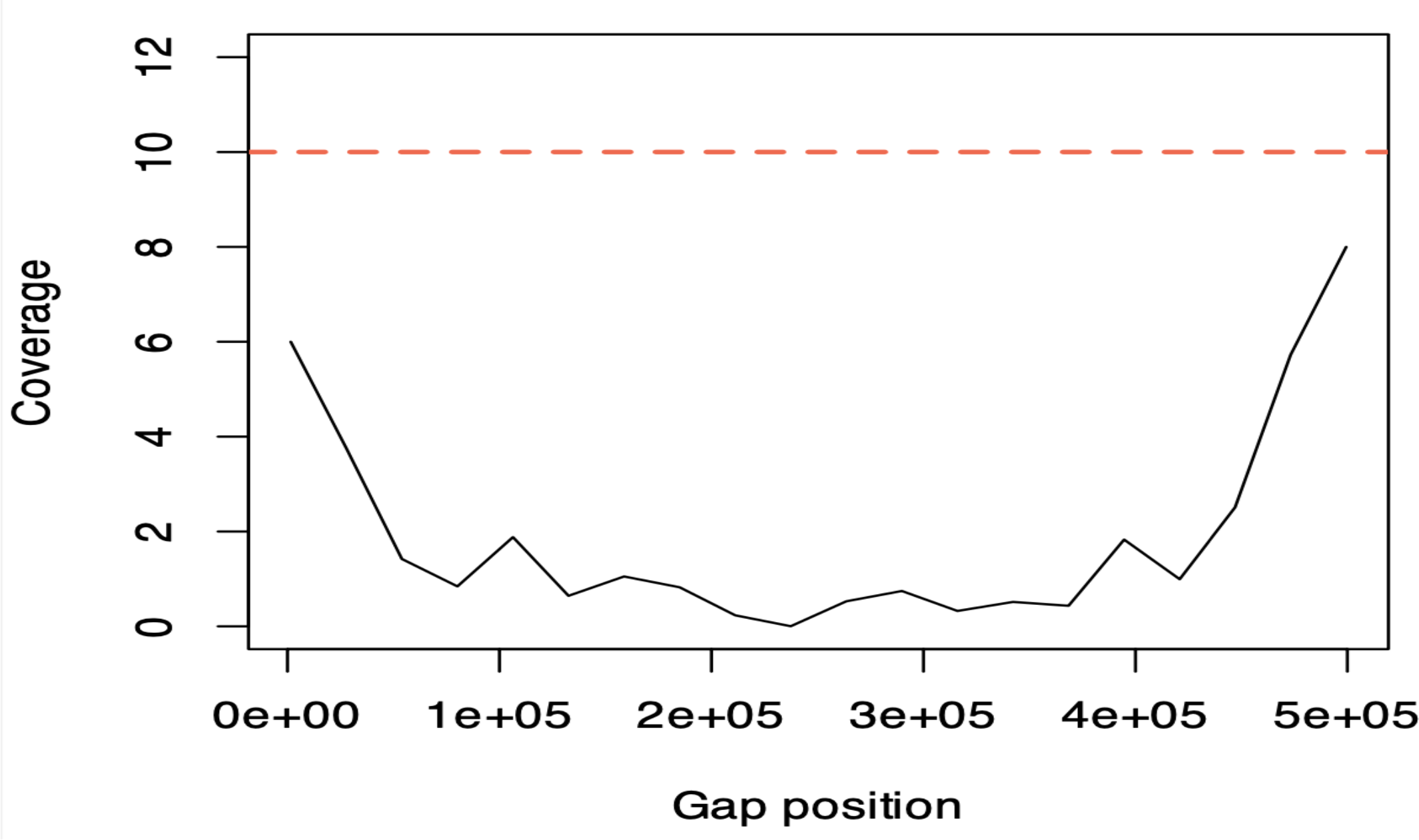
The ability of adaptive sampling to read into genome assembly gaps. Shown is the approximate sequencing depth across the relative coordinates of a 500 kbp gap constructed in hg38 chr1. The red line is the median depth of 500 kbp upstream and downstream of this gap.

There was concern that adaptive sampling may reduce the lifespan of the pores, however by using the metadata attached to each read we were able to compare the number of active channels over time in the split flow cell experiment. We observed that adaptive sampling had no obvious effect, as the number of channels decreased at the same rate for both channels performing AS and standard sequencing (Fig 6.).

**Fig 6.**
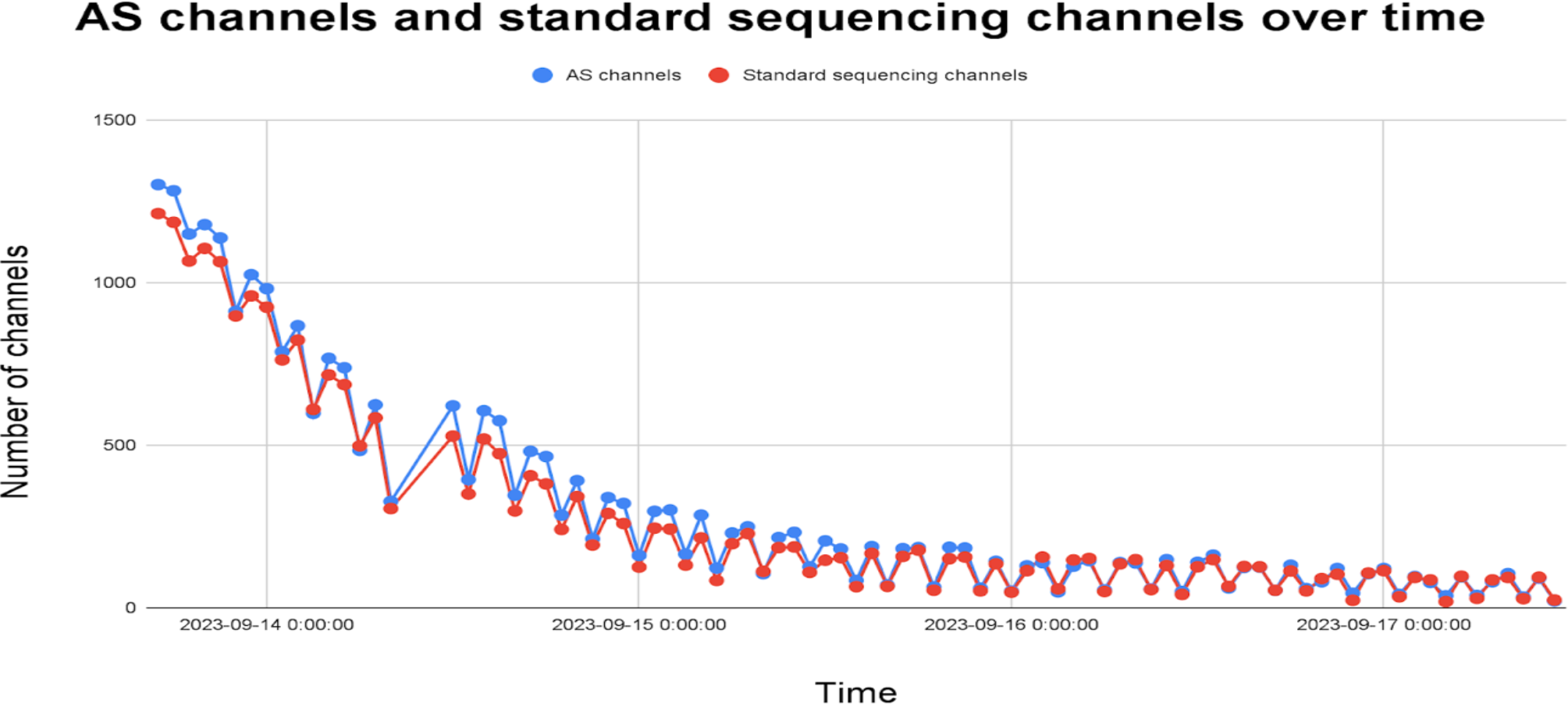
Adaptive sampling does not increase the rate of pore death. Number of available channels was measured by the PromethION over the course of the sequencing run for both channels used for adaptive sampling and standard sequencing. These varied over time, trending lower as the experiment progressed

## Conclusion

1. The coverage of ROIs is greater with adaptive sampling than with standard sequencing, with 3.6 times more coverage seen in the reserved channel flow cell test.
2. The position of the ROI within the genome has a very significant effect on the enrichment seen
3. ROI enrichment does not trend with ROI length – 10 short ROIs have the same overall enrichment as a single ROI of equivalent size.
4. Adaptive sampling does not noticeably reduce the lifespan of the ONT flow cells compared to standard sequencing.
5. Gap-filling can be performed to an extent using adaptive sampling
6. A mapping variant using the ROI as the reference genome gave a lower sequence enrichment than standard mapping (4.2x vs 6x)

Adaptive sampling does increase the coverage of ROIs relative to standard sequencing. Given the minimal additional requirements for performing adaptive sampling, it is likely to be of benefit to many researchers wishing to optimise the sequencing depth of specific target regions rather than investigating the entire genome. The flexibility of ROI length and number lends versatility to the technique, though care must be taken when designing ROIs given the strong influence position has been observed to have on relative enrichment, for example when centromeres are included. When performing such investigations the standard mapping procedure recommended by ONT should be taken as current best practice.

In addition, the “Gap-filling” capability offers a valuable area of future exploration. When coupled with the long individual strand reads of ONT sequencing it gives the potential to investigate regions with high local variance unavailable from short read sequencing techniques.

## Methods

The sequencing library used for both experiments were prepared from commercially sourced human DNA (Promega CAT#G1471) with a ligation sequencing kit V14 (ONT CAT#SQK-LSK114) following the manufacturer’s instructions. The DNA was fragmented using a g-Tube (Covaris CAT#520079) with a 50 µL volume at 1500 RCF. Final library concentration was determined to be 25.2 ng/µL via Qubit HS dsDNA (ThermoFisher CAT# Q32851) and the average size of the library was measured at 44.66 Kbp using the 5200 Fragment Analyzer System (Agilent), with an HS large fragment kit 50kb (Agilent CAT#DNF-464-0500). The trace revealed a large, normally distributed peak spanning 20-40 Kbp. The libraries were sequenced using FLO-PRO114M: R.10.4.1 flow cells (ONT) with a loading concentration of 10 fmoles (note at the time this was the recommended loading concentration, but this has since increased to 50 fmoles). Sequencing was performed on the PromethION 24 (ONT) using high-accuracy base calling model (400 bps), with the following software versions for experiment 1 and experiment 2 respectively: MinKNOW 23.07.8 and 23.04.6, Guppy 7.0.9 and 6.5.7. Run time for the majority of samples was one hour in the “multiple run” experiment and 72 hours for the “reserved channel” experiment. For the sample testing gap-filling in the “multiple run” experiment the run was extended to 19 hours as this was the final sample tested and the number of available pores had been greatly reduced when sequencing for this sample began. The control runs performed without AS to confirm the functionality of the flow cell at the beginning and halfway point of the “multiple run” experiment were run for 30 minutes.

A number of BED files targeting GRGh38 were generated, along with one custom fasta file.

1. One contiguous as possible ROI targeting 0.1, 0.25, 0.5, 1, 4 and 8% of the human genome. Made up from chr1 and chr2, excluding telomeres and centromeres. (files named “day1_0.1pct”, “day1_0.25”, etc.)
2. One bed file containing 10 ROIs at 0.1% of genome size each and one containing 80 ROIs at 0.1% each (files named “1.bed” and “2.bed”). The location of these are random, but exclude telomeres, centromeres, etc. found in Dellys’ exclude template for hg38 (https://github.com/dellytools/delly).
3. BED file targeting 0.5% (16 Mbp) of the genome with a custom hg38 assembly in which the centre 500Kbp has been masked (substituted with ‘N’ bases) to simulate an assembly gap.
4. The aforementioned 1% ROI however with a trimmed down fasta file only containing one contig, the ROI sequence (file named ROI_as_ref)

All BED file manipulations were performed using BEDTools v. 2.31.0 (Quinlan & Hall 2010). For coverage calculations, the “fastq_pass” reads were mapped to hg38 using minimap2 v. 2.26 (Heng 2018), then measured using qualimap bamQC v. 2.2.2d (Okonechnikov 2016) with the “-gff” and “-os” options. Coverage tracks were generated using deeptools bamCoverage v. 3.5.4 (Ramírez *et al*. 2016). And finally nanoplot v. 1.42.0 (De Coster 2023) generated read length histograms and read quality metrics.

## Data availability

BED files and analysis scripts can be found in the github repository, https://github.com/NationalGenomicsInfrastructure/NGI-AS-tech_note-2024

Fastq files of the basecalled called reads are submitted to ENA (accession: PRJEB73272)

## Appendix 1

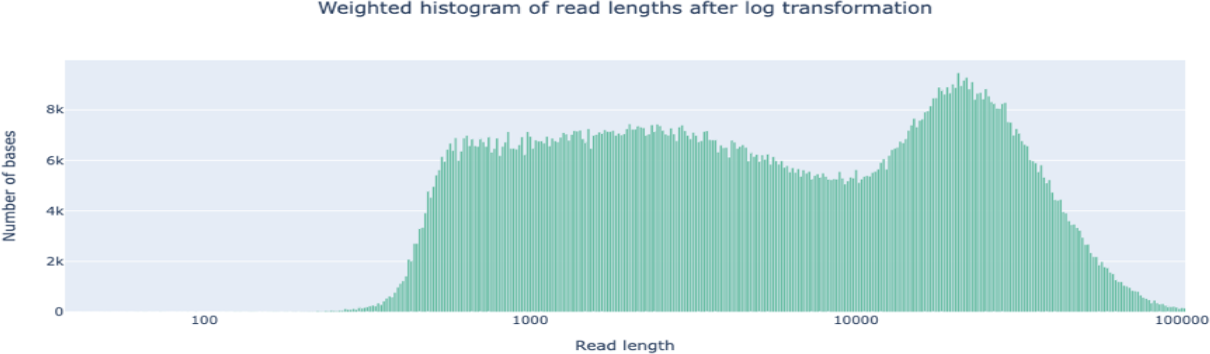

**Weighted histogram of read length with AS**

\# passed reads < 1Kb = 5,165,356

Read len N50 = 23949 (+/- 12,971.4)

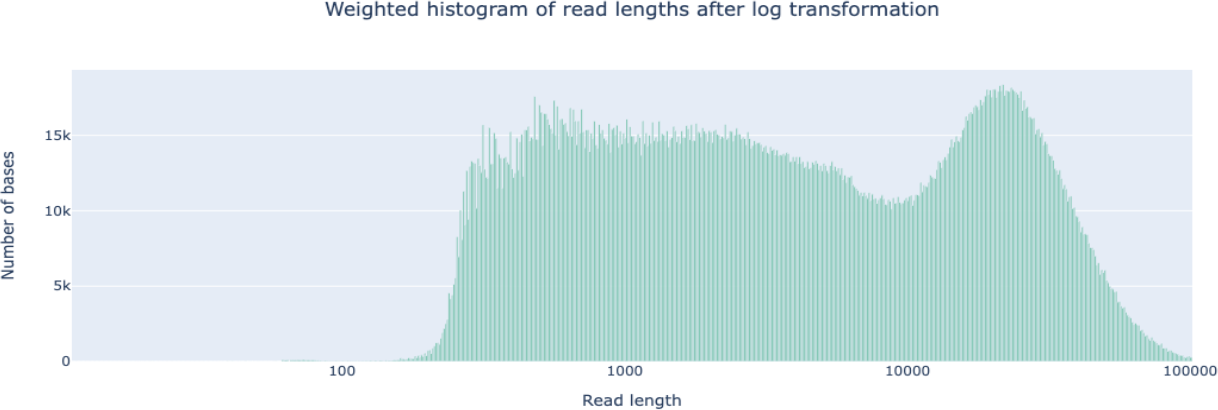

**Weighted histogram of read length with standard sequencing**

\# passed reads < 1Kb = 574,268

Read len N50 = 23647 (+/- 12,241.1)

## References

https://community.nanoporetech.com/info_sheets/adaptive-sampling/v/ads_s1016_v1_revj_12nov2020/considerations-for-experimental-design

Wouter De Coster, Rosa Rademakers, NanoPack2: population-scale evaluation of long-read sequencing data, Bioinformatics, Volume 39, Issue 5 (2023) btad311, 10.1093/bioinformatics/btad311

Starke, H., Seidel, J., Henn, W. et al. Homologous sequences at human chromosome 9 bands p12 and q13-21.1 are involved in different patterns of pericentric rearrangements. Eur J Hum Genet 10, 790–800 (2002). 10.1038/sj.ejhg.5200889

Li, Heng. “Minimap2: pairwise alignment for nucleotide sequences.” Bioinformatics 34.18 (2018): 3094–3100.

Martin, S., Heavens, D., Lan, Y. et al. Nanopore adaptive sampling: a tool for enrichment of low abundance species in metagenomic samples. Genome Biol 23, 11 (2022). 10.1186/s13059-021-02582-x

Konstantin Okonechnikov, Ana Conesa, Fernando García-Alcalde, Qualimap 2: advanced multi-sample quality control for high-throughput sequencing data, Bioinformatics, Volume 32, Issue 2 Pages 292–294 (2016) 10.1093/bioinformatics/btv566

Ramírez, Fidel, Devon P. Ryan, Björn Grüning, Vivek Bhardwaj, Fabian Kilpert, Andreas S. Richter, Steffen Heyne, Friederike Dündar, and Thomas Manke. deepTools2: A next Generation Web Server for Deep-Sequencing Data Analysis. Nucleic Acids Research (2016). doi:10.1093/nar/gkw257.

Quinlan AR and Hall IM BEDTools: a flexible suite of utilities for comparing genomic features. Bioinformatics. 26, 6, pp. 841–842 (2010). 10.1093/bioinformatics/btq033

Weilguny, L., De Maio, N., Munro, R. et al. Dynamic, adaptive sampling during nanopore sequencing using Bayesian experimental design. Nat Biotechnol 41, 1018– 1025 (2023). 10.1038/s41587-022-01580-z

